# Notes on *f*_4_-ratio estimation

**DOI:** 10.64898/2025.12.22.695918

**Authors:** Kalle Leppälä

## Abstract

The *f*_4_*-ratio estimation* is a popular technique for estimating gene flow proportion as a ratio between two *f*_4_-statistics. We point out the underappreciated fact that the root is not important in *f*_4_-ratio estimation, making the technique highly versatile. We study robustness and the bidirectional case. We introduce an *enhanced* branch statistic *f*_*B*_ for detecting tentative gene flow events in a large phylogenetic tree, which has favourable properties compared to Malinsky’s branch statistic *f*_*b*_.

## Introduction

The *f*_4_*-ratio estimation* is a popular technique for estimating gene flow proportion as a ratio between two *f*_4_-statistics in certain admixture graphs. Its first use as well as the first use of *f*_4_-statistics themselves was by Reich et al. [35], where the technique was called *f*_4_ *ancestry estimation*. It was re-introduced formally as *f*_4_-ratio estimation by Patterson et al. [29]. Both articles assume the rooted admixture graph marked with an asterisk (∗) in Figure 1. The root is however not actually needed for anything. At the center of Figure 1 is an unrooted five-leaf admixture graph that can be rooted in five different ways (ignoring labels) shown in the inner ring. The outer ring shifts perspective further, where one admixture branch is seen as an integral part of a *backbone* tree, and the other one supplementing the tree with a gene flow event. This is the framework behind some popular gene flow detection methods such as the *D*-statistic [12, 29] and the branch statistic *f*_*b*_ [26, 25]. In (supplementary) Figure S11.2 Palkopoulou et al. [28] apply the *f*_4_-ratio estimation without a root in a situation corresponding to the admixture graph marked with a dagger (†). Hence, the fact that the admixture proportion in all gene flow events visible in Figure 1 can be estimated using *f*_4_-ratios is not necessarily novel information, but we suggest it is not widely recognized. This work is an attempt to highlight the versatility of *f*_4_-ratio estimation (being applicable to a total of nine distinct gene flow scenarios) to a wider audience. We derive formulae for bidirectional gene flow. We adapt the lessons for the betterment of the branch statistic *f*_*b*_, introducing an *enhanced* statistic *f*_*B*_ that has in many ways preferable behaviour. Finally, we apply both the *f*_4_-ratio estimation and and the enhanced branch statistic *f*_*B*_ to real world genetic data and correct some misunderstandings about gene flow between brown bears and polar bears.

**Figure 1.**
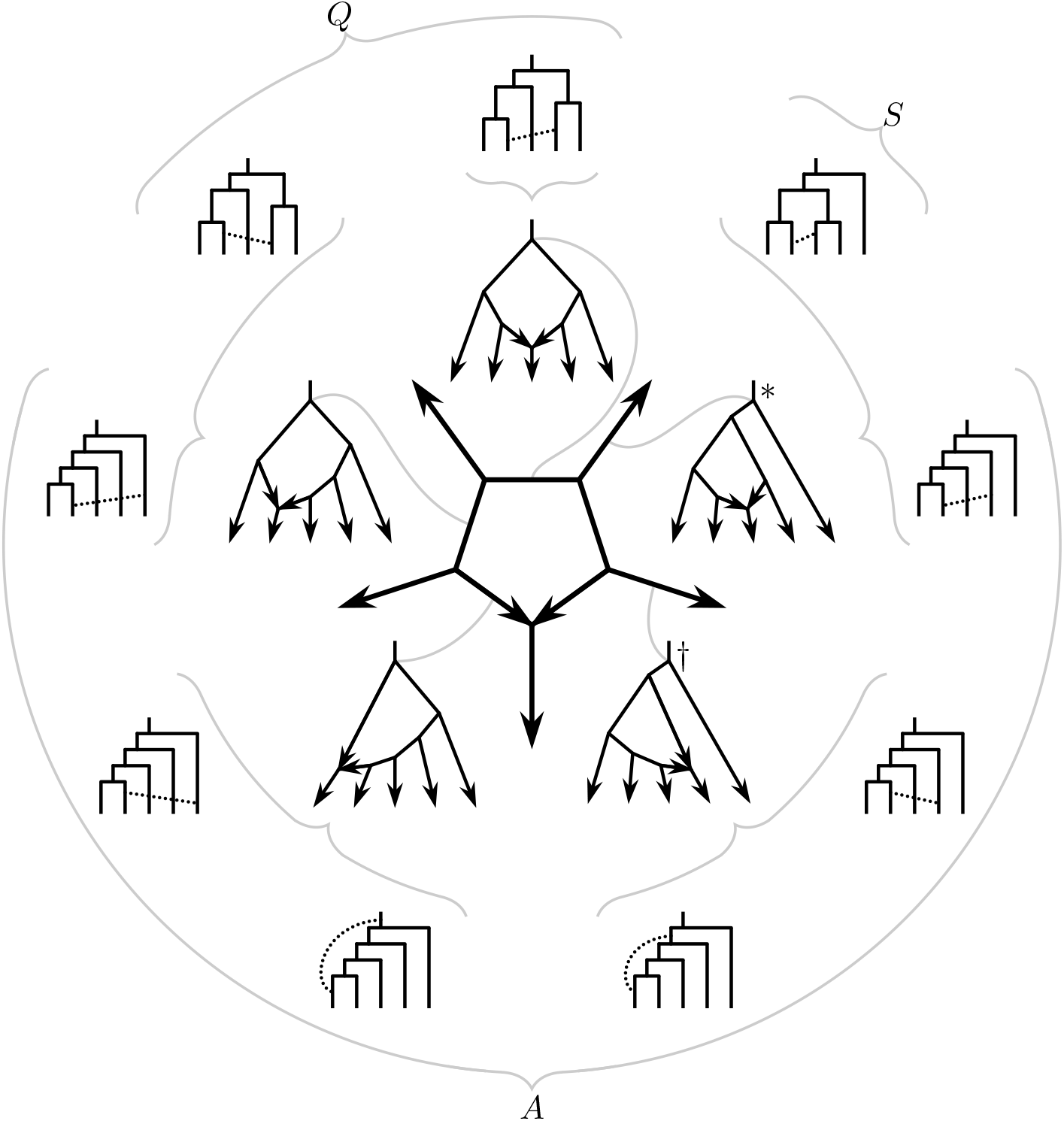
Three perspectives. The unrooted admixture graph in the middle can be rooted in five different ways (ignoring labels, note that the root cannot be placed after the admixture event), shown on the inner ring. Either one of the admixture edges could be seen as integral part of the backbone tree and the other one as supplementing admixture event. The nine scenarios obtained this way are depicted on the outer ring. These are precisely the nine gene flow events that can be uniquely identified using Δ-statistics [21] without singleton patterns, in other words, using *D*-statistics [12, 29] repeatedly.

## General *f*_4_-ratio estimation

The *f*_4_-statistic *f*_4_(*W, X*; *Y, Z*) in an admixture graph where branch lengths measure *units of drift* can be seen as the directed overlap between admixture proportion weighted paths from *W* to *X* and from *Y* to *Z* [29]. In short, a path is allowed to go first upstream and then downstream (but never upstream again after switching), and it is allowed to bifurcate and rejoin, acquiring segment specific weights from admixture branches while split. The overlap between paths traveling opposite directions is counted negative.

Figure 2 left panel visualizes the method of *f*_4_-ratio estimation. Notation varies between sources — already the identities

**Figure 2.**
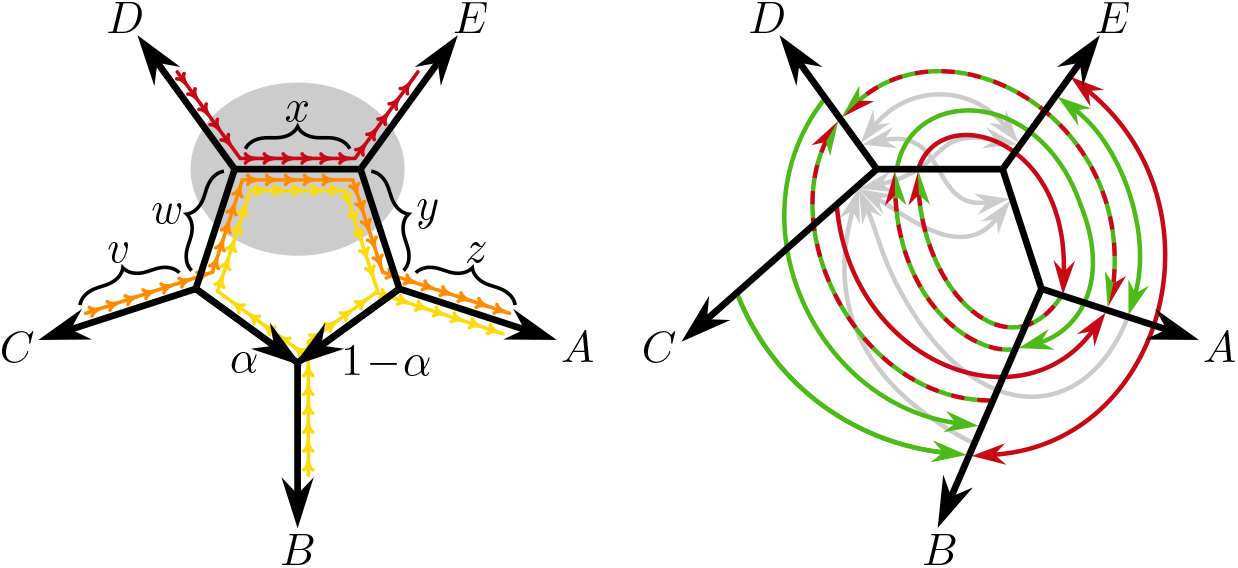
Left: the general visual logic behind *f*_4_-ratio estimation. Right: how the ratio *f*_4_(*B, A*; *D, E*)*/f*_4_(*C, A*; *D, E*) is influenced by gene flow events other than *C* → *B* (green positively, red negatively, striped either way depending on graph parameters, grey neutral). Note that the double headed arrow in this figure is only a space saving device — it does not stand for bidirectional gene flow.

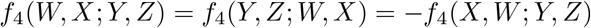

inflict considerable ambiguity. We have adapted our own notation to stress that no leaf is considered an outgroup; *B* is the hy(B)rid, *C* is the sour(C)e of gene flow of proportion *α* we’re interested in, and *A, D* and *E* then fall into place.

The *f*_4_-ratio estimate of *α* is simply

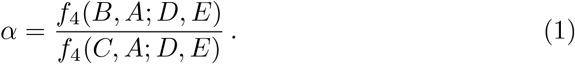

The yellow path from *B* to *A*, the orange path from *C* to *A* and the red path from *D* to *E* all traverse first upstream and then downstream no matter where the root is placed (it cannot be place on the terminal branch leading to leaf *B*). The yellow path is split while in the pentagon, being equipped with weight *α* on the detour and weight 1 − *α* on the shortcut. Therefore *f*_4_(*B, A*; *D, E*) = *αx* as a weighted overlap between the yellow path and the red path, and *f*_4_(*C, A*; *D, E*) = *x* as a weighed overlap between the orange path and the red path, and (1) is shown. Estimating 1 − *α* instead of *α* is just a matter or relabeling. If only four leaves are available, compromises bounding the admixture proportion from below and above are easily derived from Figure 2 by replacing the red path with either the path from *C* to *E* or with the path from *D* to *A*:

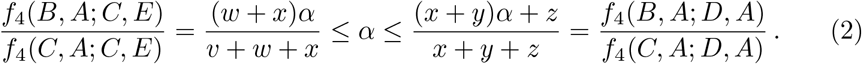

### Robustness

In order for the interpretation of (1) as an admixture proportion to be justified, the admixture graph must satisfy the assumptions made. For example, Peter [31] points out a common pitfall in Figure 6 panel C: in a tree ((((*D, C*), *B*), *A*), *E*) the proportion (1) lies between zero and one without any gene flow at all. This is because the tree has the wrong shape — it does not match the model in Figure 2 left panel even if we were to remove one of the two admixture edges.

But we need not be overtly strict. Inside the grey ellipse in Figure 2 left panel, the yellow and the orange paths travel together with the only exception that the weight of the yellow path is *α* while the weight of the orange path is one. This means that after adding any amount of admixture edges inside the grey ellipse, the yellow and orange paths will travel the same bifurcating and rejoining route, however complicated, only differing on their weights everywhere by the factor *α*. It follows that whatever they have in common with the red path — here *x* — also differ only by the factor *α* and that (1) is unaffected.

Sometimes *f*_4_-ratio estimation is used as a means to detect a directed gene flow event rather than estimate the gene flow proportion of an already established event. This approach has the caveat that *C* → *B* with *α >* 0 is only one possible explanation for an observation of *f*_4_(*B, A*; *D, E*)*/f*_4_(*C, A*; *D, E*) *>* 0 even if the backbone tree shape of the phylogeny is assumed. Figure 2 right panel visualizes the effect of all 24 possible gene flow events between non-neighbouring edges on the *f*_4_-ratio. Green arrow indicates that the ratio will be positive, red negative, striped either positive or negative depending on graph parameters, and grey zero. Here the double headed arrows are a space saving notation for two arrows. As the ratio is a composition of two *f*_4_-statistics, the situation is necessarily quite complex, but some features deserve highlighting. First, incoming arrows to branch *C* are all grey because gene flow into an unsampled population can’t be detected, and from point of view of the numerator *f*_4_(*B, A*; *D, E*) the leaf *C* is unsampled. Second, even if *B* truly is the target of gene flow, the source could also be *D* or the inner edge *C, D* | *E, A, B*. Note that while the argument about the grey ellipse on the left panel allows unlimited number of admixture edges, the effects on the *f*_4_-ratio shown on the right panel are for strictly one admixture event. If the admixture proportions are low, and paths traversing through several different gene flow events therefore negligible, then the effects are roughly additive. If not, already *B* → *C* and subsequent *C* → *D* — two zero effect admixture events — can produce a positive numerator *f*_4_(*B, A*; *D, E*) and a denominator *f*_4_(*C, A*; *D, E*) of any sign.

## Bidirectionality

As seen in Figure 2 right panel, gene flow of the reverse direction *B* → *C* cannot alone explain a positive *f*_4_-ratio estimator. However, the arrow being grey does not mean that it does not interfere in any way. Already in Supplementary Figure 19 of [20] we pointed out that if the gene flow 2 → 3 in admixture graph marked with an asterisk (∗) in Figure 1 is actually bidirectional with *β* denoting the admixture proportion of the opposite direction 3 → 2 (Figure 3), then the expected value of the *f*_4_-ratio estimator is *α/*(1 − *β*). We recognize that when gene flow is low, the bidirectionality bias is negligible, but with high levels of admixture the unidirectional estimates could up to double the correct value. Here we repeat the argument of Supplementary Figure 19 of [20] in full generality that applies to four bidirectional gene flow settings, and solve the unbiased admixture proportion estimators.

**Figure 3.**
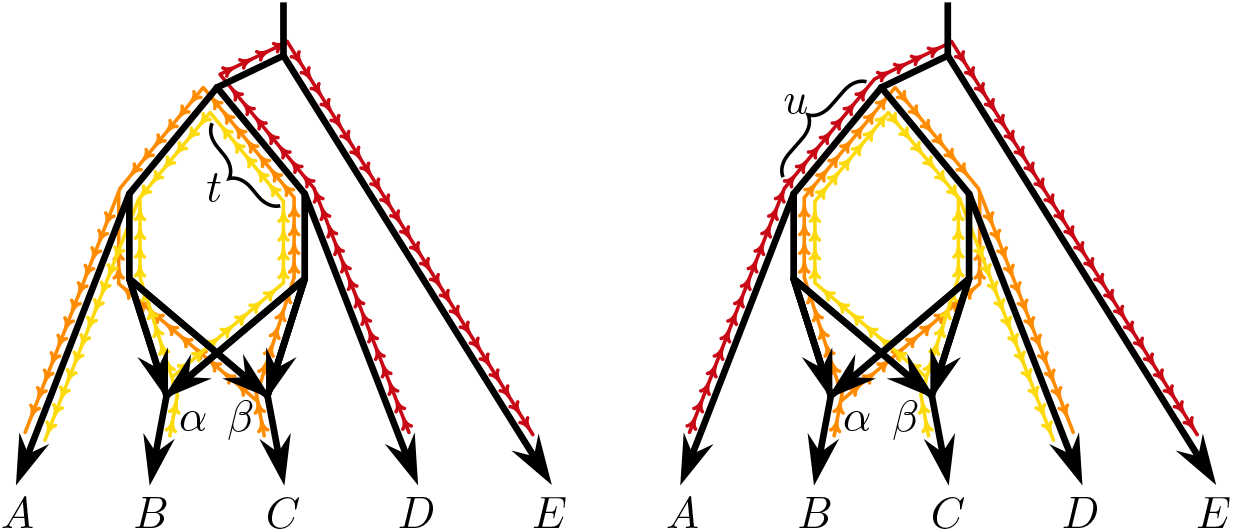
Left: Estimating the proportion of gene flow 3 → 2 using the *f*_4_-ratio gives *f*_4_(*B, A*; *D, E*)*/f*_4_(*C, A*; *D, E*) = *αt/*((1 − *β*)*t*) = *α/*(1 − *β*). Right: When estimating the proportion of the opposite gene flow 2 → 3, the leaves *A, B, C* and *D* need to switch roles because *C* is now the target of gene flow and so on. We get *f*_4_(*C, D*; *A, E*)*/f*_4_(*B, D*; *A, E*) = *βu/*((1 − *α*)*u*) = *β/*(1 − *α*). Both estimators are biased, but unbiased estimators (4) are readily solved (3).

In the bidirectional case, the numerator of (1) (the overlap between the yellow and red paths in Figure 2 left panel) is unaffected, but the denominator (the overlap between the orange and red paths) attains the weight 1 − *β*. This is the first identity in (3). Switching the roles of *α* and *β*, see Figure 3, yields the second identity, and unbiased estimators (4) of *α* and *β* are solved from the obtained linear system. We remark that potential gene flow events inside the grey ellipse in the left panel of Figure 2 are no longer harmless after switching the roles of *α* and *β*, in other words, that bidirectional admixture proportion estimation requires stricter assumptions than the usual *f*_4_-ratio does.

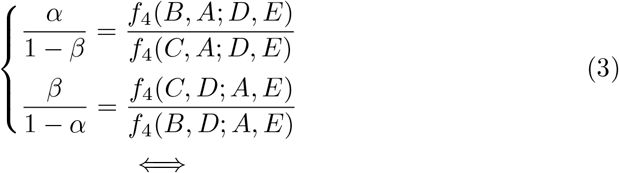

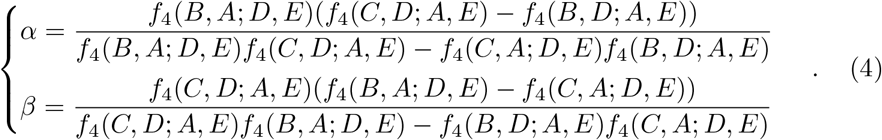

This unrooted reasoning applies in a total of four instances up to relabeling of leaves: gene flow between 2 and 3 in tree *S* = (((1, 2), (3, 4)), 5) (Figure 3), gene flow between 2 and 4 or between 2 and 5 in tree *A* = ((((1, 2), 3), 4), 5), and gene flow between 2 and 5 in tree *Q* = (((1, 2), 3), (4, 5)). The explicit formulae are listed in Table 1.

**Table 1.** Bidirectional gene flow estimators in the four special cases the rootless reasoning applies to.

| Tree | Gene flow | Bidirectional estimates |
| --- | --- | --- |
| $S$ | $\alpha: 3 \rightarrow 2$ | $\frac{f_4(2, 1; 4, 5)(f_4(3, 4; 1, 5) + f_4(2, 4; 1, 5))}{f_4(2, 1; 4, 5)f_4(3, 4; 1, 5) - f_4(3, 1; 4, 5)f_4(2, 4; 1, 5)}$ |
| | $\beta: 2 \rightarrow 3$ | $\frac{f_4(3, 4; 1, 5)(f_4(2, 1; 4, 5) + f_4(3, 1; 4, 5))}{f_4(3, 4; 1, 5)f_4(2, 1; 4, 5) - f_4(2, 4; 1, 5)f_4(3, 1; 4, 5)}$ |
| $A$ | $\alpha: 2 \rightarrow 4$ | $\frac{f_4(4, 5; 1, 3)(f_4(2, 1; 5, 3) - f_4(4, 1; 5, 3))}{f_4(4, 5; 1, 3)f_4(2, 1; 5, 3) - f_4(2, 5; 1, 3)f_4(4, 1; 5, 3)}$ |
| | $\beta: 4 \rightarrow 2$ | $\frac{f_4(2, 1; 5, 3)(f_4(4, 5; 1, 3) - f_4(2, 5; 1, 3))}{f_4(2, 1; 5, 3)f_4(4, 5; 1, 3) - f_4(4, 1; 5, 3)f_4(2, 5; 1, 3)}$ |
| $A$ | $\alpha: 2 \rightarrow 5$ | $\frac{f_4(5, 4; 1, 3)(f_4(2, 1; 4, 3) - f_4(5, 1; 4, 3))}{f_4(5, 4; 1, 3)f_4(2, 1; 4, 3) - f_4(2, 4; 1, 3)f_4(5, 1; 4, 3)}$ |
| | $\beta: 5 \rightarrow 2$ | $\frac{f_4(2, 1; 4, 3)(f_4(5, 4; 1, 3) - f_4(2, 4; 1, 3))}{f_4(2, 1; 4, 3)f_4(5, 4; 1, 3) - f_4(5, 1; 4, 3)f_4(2, 4; 1, 3)}$ |
| $Q$ | $\alpha: 2 \rightarrow 4$ | $\frac{f_4(4, 5; 1, 3)(f_4(2, 1; 5, 3) - f_4(4, 1; 5, 3))}{f_4(4, 5; 1, 3)f_4(2, 1; 5, 3) - f_4(2, 5; 1, 3)f_4(4, 1; 5, 3)}$ |
| | $\beta: 4 \rightarrow 2$ | $\frac{f_4(2, 1; 5, 3)(f_4(4, 5; 1, 3) - f_4(2, 5; 1, 3))}{f_4(2, 1; 5, 3)f_4(4, 5; 1, 3) - f_4(4, 1; 5, 3)f_4(2, 5; 1, 3)}$ |

### Validating the bidirectional estimators with a simulation

To validate the proposed estimators, we simulated tree *S* = (((1, 2), (3, 4)), 5) with gene flow between 2 and 3 of proportions 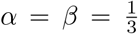. We used *SLiM* [15] to (forward) simulate the history after the root without any mutations, and *msprime* [3] to (coalescent) simulate the history before the root and adding neutral mutations. All populations at all times consisted of 10,000 individuals with diploid genomes of two times 10 chromosomes of length 10^8^. The mutation rate and the recombination rate were both 10^−8^, per site per generation. The instantaneous admixture event happened 10,000 generations ago, and the populations 1 and 2, 3 and 4, 12 and 34, and 1234 and 5 splitted 20,000, 20,000, 40,000 and 60,000 generations ago, respectively.

The gene flow proportions were then estimated with *admixtools* [24] and R using the biased estimator (1) and the unbiased estimators (4). Standard errors were constructed with the block jackknife procedure (the *δ*-method at times returned negative estimators of variance). Naïvely applying (1) without correcting for bias yielded *α* = 0.507 (SE = 0.006) and *β* = 0.504 (SE = 0.006), which was to be expected as 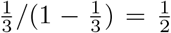. Being conscious of the possible bidirectional nature of the admixture event and using (4) yielded *α* = 0.338 (SE = 0.004) and *β* = 0.334 (SE = 0.004), consistent with the true values.

All the code is stored at https://github.com/KalleLeppala/rootless.

## Enhanced branch statistics *f*_*B*_

Malinsky’s branch statistic *f*_*b*_ [26, 25] aims to summarize excess allele sharing between a terminal branch *s* and another (possibly inner) branch *t*, in order to infer putative gene flow events in a large phylogenetic tree. The *f*_*b*_-statistic is essentially *f*_4_-ratio estimation using the conservative compromise (2) (first inequality), applied on all subtrees that include the outgroup. With our unrooted perspective on *f*_4_-ratio estimation, we immediately recognize that an outgroup is not really necessary. In fact, it turns out that requiring the use of outgroup is actively harmful, as is forcing the compromise (2). In this section we describe the *enhanced* branch statistic *f*_*B*_, give theoretical motivations behind the altered definition, and demonstrate using simulations that *f*_*B*_ has desirable properties over *f*_*b*_.

In a tree, any two branches *s* and *t* are connected by a unique sequence of branches *p*_1_, *p*_2_, …, *p*_*n*_. In accordance to the diagram in top left panel of Figure 4, we define 𝒞 as the set of leaves on the side of *s* that *t* is not, 𝒜 as the set of leaves on the side of *t* that *s* is not, and 𝒮_*i*_ as the set of leaves that equally far from *p*_*i*−1_ and *p*_*i*_ — here we think of *s* as *p*_0_ and of *t* as *p*_*n*+1_. Now if 𝒟 = *S*_*i*_ and ℰ = *S*_*j*_, *i < j*, and *A, B, C, D, E* are any leaves from their respective sets 𝒜, ℬ, 𝒞, 𝒟, ℰ, we can estimate the proportion *α* of the gene flow from *s* into *t* as (1), see left panel in Figure 2. Furthermore, with 𝒟 = 𝒞 the left hand side proportion (2) is a lower bound for *α*. We make the convention 𝒮_0_ = 𝒞 and are ready to set

**Figure 4.**
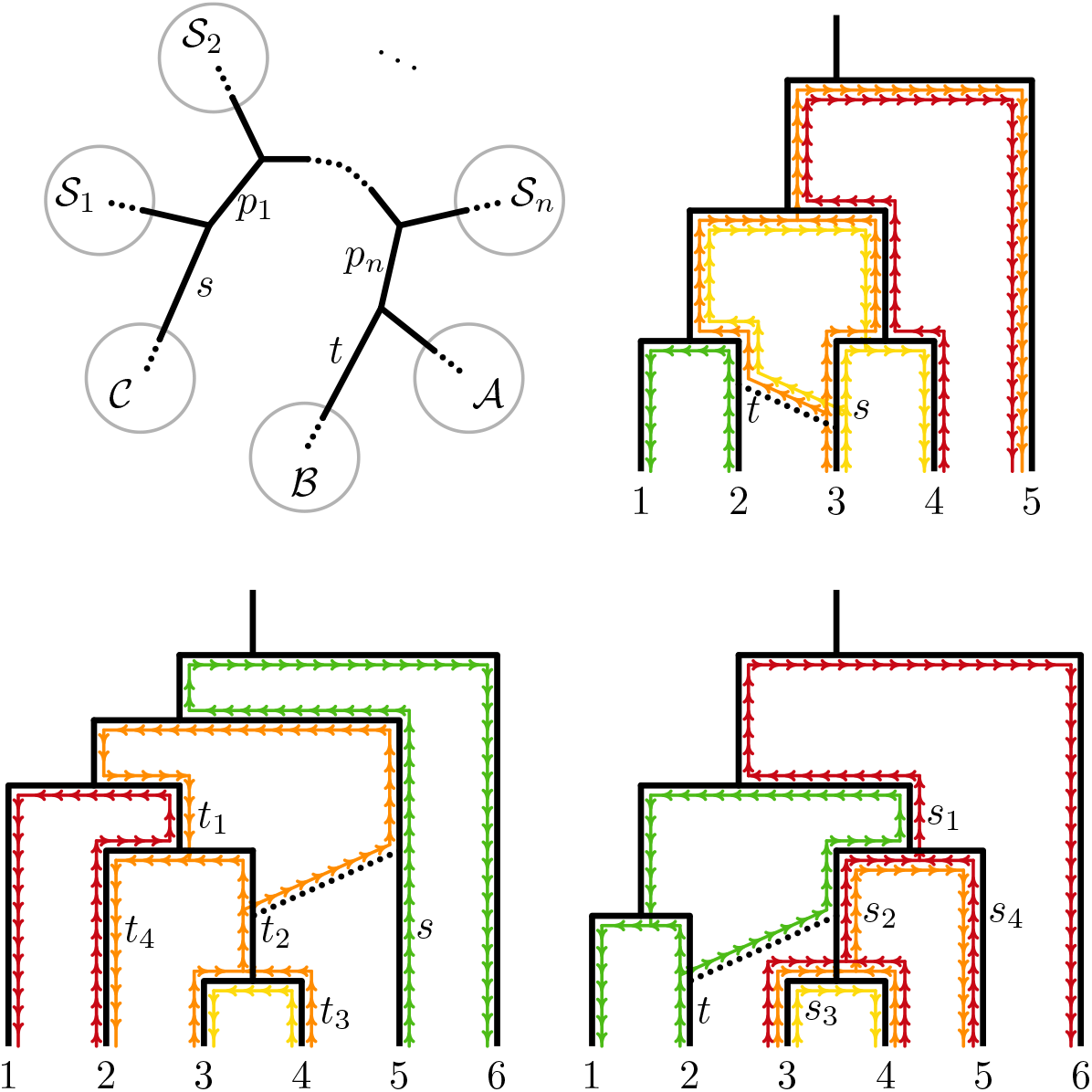
Top left: the setup for the enhanced *f*_*B*_-statistic testing for gene flow *s* → *t*. Top right: a situation where Malinsky’s *f*_*b*_ gives a false positive signal. Bottom left: identifying the target of gene flow. Bottom right: identifying the source of gene flow. For full explanations, see the main text.

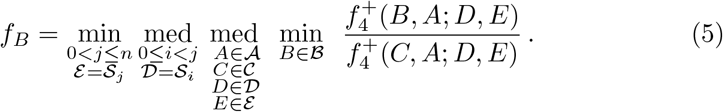

This definition warrants some clarifications. The operator med stands for the *lower median* that attains the lower of the middle two values when the set has even cardinality. The plus in the *f*_4_-statistic is shorthand for

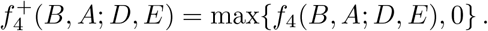

The fraction is not defined unless *f*_4_(*C, A*; *D, E*) is strictly positive, in practice statistically significant using a one-tailed test. Operators min and med are to be understood as taken over only those fractions that exist. Now, *f*_*B*_ is a non-negative real number if defined, and a positive value is interpreted as tentative presence of gene flow *s* → *t* rather than literal admixture proportion.

Malinsky’s *f*_*b*_ differs from definition (5) in several ways: *i* = 0 so that 𝒟 = 𝒞, *s* is assumed to be a terminal branch so that | 𝒞 | = 1, and the outgroup alone serves the role of ℰ. Next we motivate the enhanced definition using theoretical examples illustrated in Figure 4. The reasoning is meant to be heuristic rather than conclusive. As both *f*_*b*_ and *f*_*B*_ ultimately rely on repeated application of the *f*_4_-ratio estimation without paying much mind to the required assumptions, some amount of error is unavoidable.

### Identifying the target of gene flow

Figure 4 bottom left panel depicts a tree equipped with a gene flow event *s* → *t*_2_. Ideally a statistic does not confuse the scenario with gene flow into an ancestor branch *t*_1_, a descendant branch *t*_3_, or a sister branch *t*_4_. Testing for gene flow *s* → *t*_1_ uses

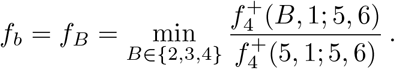

Because 2 is not affected by gene flow, the red path and the green path do not overlap, and therefore *f*_4_(2, 1; 5, 6) = *f*_*b*_ = *f*_*B*_ = 0. We can appreciate how taking a minimum over the set ℬ instead of a median in definitions of *f*_*b*_ and *f*_*B*_ was the key in recognizing that *s* → *t*_1_ does not happen. Testing for (the correct) gene flow *s* → *t*_2_ yields

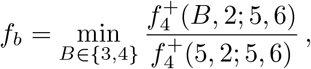

so Malinsky’s *f*_*b*_ straightforwardly gives a positive result as the orange path and the green path overlap no matter which starting leaf 3 or 4 we use for the orange path. The enhanced statistic

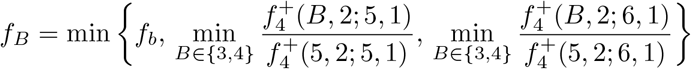

remains positive because the paths from 6 or 5 to 1 (not drawn) also positively overlap with the orange paths. Testing for gene flow *s* → *t*_3_ will not detect a signal;

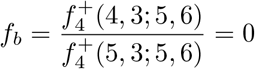

because the yellow path and the green path do not overlap, and *f*_*B*_ = 0 as a minimum of a set that contains *f*_*b*_. Finally, testing for *s* → *t*_4_ is done with

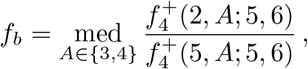

which is zero, because *f*_4_(2, *A*; 5, 6) is negative as can be seen by observing the overlap between the green path and the reversed orange path, whichever leaf it ends at. The enhanced statistic *f*_*B*_ is a minimum of a set that contains *f*_*b*_ and is then also zero. Hence, both statistics *f*_*b*_ and *f*_*B*_ could correctly identify the target of gene flow. For Malinsky’s *f*_*b*_, comparable reasoning was already presented in [25], and is only repeated here for completeness’ sake.

### Identifying the source of gene flow

The tree in Figure 4 bottom right panel has a gene flow event *s*_2_ → *t*. We wish to be able to identify the source branch *s*_2_ without mixing it up with an ancestor branch *s*_1_, a descendant branch *s*_3_, or a sister branch *s*_4_. Note that strictly speaking Malinsky’s *f*_*b*_ is not defined when the source is an inner branch (when |𝒞| *>* 1), but here we nevertheless treat it as if it was, taking a median over the set 𝒞. With this convention, *s*_1_ → *t* is tested with

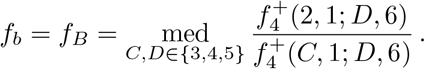

The green path and the red path overlap no matter what starting leaf 3, 4 or 5 we use for the red path, and so both statistics return a false positive result.

Testing for (the correct) gene flow *s*_2_ → *t* is done with

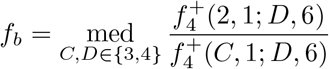

or

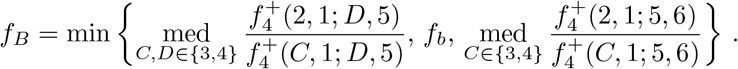

Because all the red paths and all the orange paths overlap with the green path, both statistics obtain a positive value. Malinsky’s *f*_*b*_ used to test for *s*_3_ → *t* is

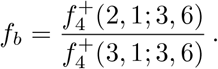

This is a positive number because the red path from 3 to 6 overlaps with the green path. The enhanced statistic *f*_*B*_ on the other hand uses many sets 𝒟 and ℰ, importantly ℰ is not forced to be the outgroup {6}. When ℰ = {4}, we must have 𝒟 = {3} and the statistic gets value zero because the yellow path and the green path do not overlap. Last, to detect *s*_4_ → *t* we use either

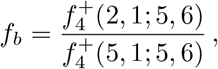

which is positive as the red path from 5 to 6 overlaps with the green path, or

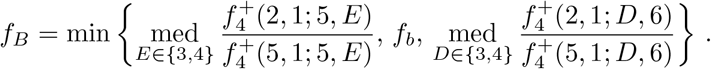

We can see that *f*_*B*_ = 0 because when 𝒟 = {5} and ℰ = {3, 4}, the statistic *f*_4_(2, 1; *D, E*) is negative as the overlap between the green path and the reversed orange path, and *f*_4_(5, 1; *D, E*) is still positive. In summary, while both statistics return a false positive signal for *s*_1_ → *t*, the enhanced statistic *f*_*B*_ could correctly reject *s*_3_ → *t* and *s*_4_ → *t* while Malinsky’s *f*_*b*_ could not. If several possible sources have positive enhanced statistic *f*_*B*_ for the same target, a careful interpretation is now believing the source where *n*, the size of the loop the gene flow event would create on the phylogenetic tree, is largest. This is not ideal because several gene flow events could be real simultaneously, but still desirable compared to *f*_*b*_.

### Reverse gene flow

The top right panel of Figure 4 illustrates another common situation which makes Malinsky’s *f*_*b*_ return a false positive. The actual gene flow is *t* → *s*, but we are testing for *s* → *t*. Now

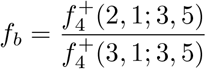

is strictly positive because the orange path and the green path overlap. The enhanced statistic *f*_*B*_ is not duped;

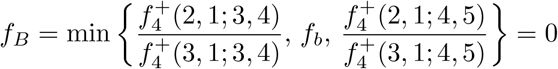

because the red path from 4 to 5 does not overlap with the green path from 2 to 1. We remark that the reason for the false positive signal was forcing the use of 𝒮 in the role of 𝒟. While that choice at first glance appears conservative, being a lower bound for the admixture proportion, it is that only when the assumptions of the *f*_4_-ratio estimation are valid which is not the case when guarding against false positives. Note that the median operator being defined as the lower median was essential here, ensuring min{*x*, med{*y, z*}} = min{*x, y, z*} (which was also applied but not needed in the previous examples). Note that even *f*_*B*_ cannot reject *s* → *t* if, say, 4 doesn’t exist and neither do the yellow and the red paths. It could be justified to require that *n* ≥ 2 in (5) to avoid the issue.

These examples illustrate why the enhanced statistic *f*_*B*_ is defined the way it is in (5). Because of the case *s* → *t*_1_ we need to take minimum over *B* ∈ ℬ. Because of the cases *s*_3_ → *t* and *s*_4_ → *t*, a zero fraction when ℰ = 𝒮_1_ needs to imply *f*_*B*_ = 0. Because of the reverse gene flow, if 𝒟 ≠ 𝒞 implies a zero fraction, it also needs to imply *f*_*B*_ = 0. A simple way to achieve these conditions would be to minimize over all sets 𝒟 and ℰ, but we wish to favor the median operator over the minimum operator to the extent it’s possible. Because of outliers, excessive minimizing could lead to false negatives especially in large phylogenies, whereas the medium operator is robust in that respect.

### Re-analysis of a simulated Malawi cichlid dataset

We re-analyzed the simulated Malawi cichlid data that were originally used to benchmark *f*_*b*_ [26, 25]. The data and the phylogenetic tree are available at https://github.com/millanek/Dsuite?tab=readme-ov-file. The results are discussed in Figure 5. We used the R package *admixtools* [24], the scripts can be found at https://github.com/KalleLeppala/rootless.

**Figure 5.**
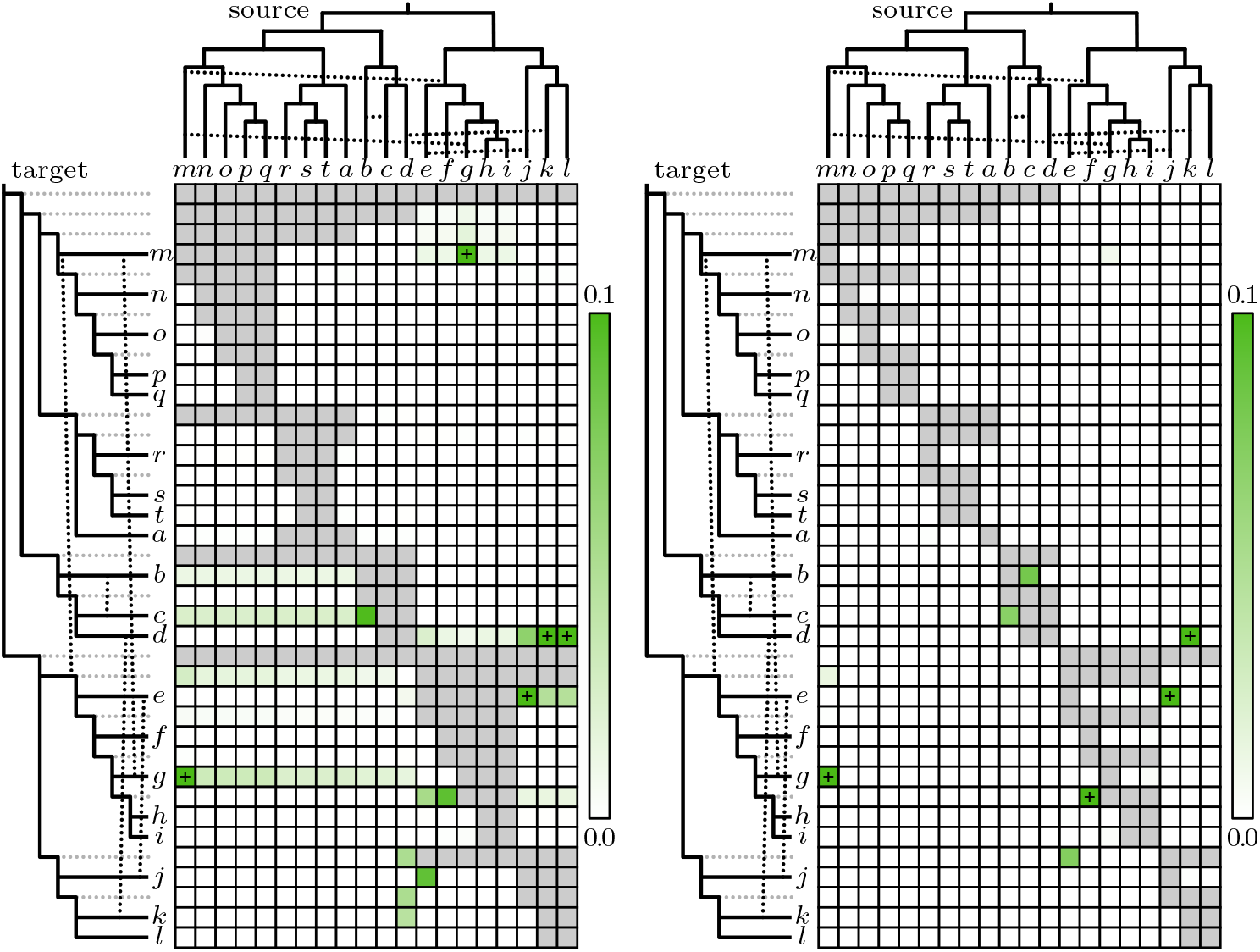
Left: reproduction of Figure 3 in [25]. Right: the same data analysed using enhanced statistics *f*_*B*_ instead. Grey cell means that the statistic is undefined. The plus symbol means that the value is over 0.1 (the highest values are 0.149 and 0.142, respectively). The branch lengths in the phylogenetic trees are not in scale, and the five gene flow events are *m* → *g* (*α* = 0.14), *m* → *efghi* (*α* = 0.08), *c* → *b* (*α* = 0.16), *k* → *d* (*α* = 0.18) and *j* → *e* (*α* = 0.13). Malinsky’s *f*_*b*_ doesn’t recognize *c* → *b* because the outgroup belongs to the set 𝒜 which is forbidden. Otherwise both methods flag all true events. The enhanced statistic *f*_*B*_ is superior in weeding out false positives, although *g* → *m, b* → *c, f* → *hi* and *e* → *jkl* remain. The first of these is inflicted by the combination of *m* → *g, m* → *efghi* and *jkl* → *e*, but the issue is less severe than it is for *f*_*b*_. The rest would be fixed by requiring *n* ≥ 2 in definition (5), but that would also reduce the range where *f*_*B*_ is defined which currently is far more generous than the range of *f*_*b*_. See Supplementary figures 1 and 2 for plots where source is allowed to be an inner branch.

**Figure 6.**
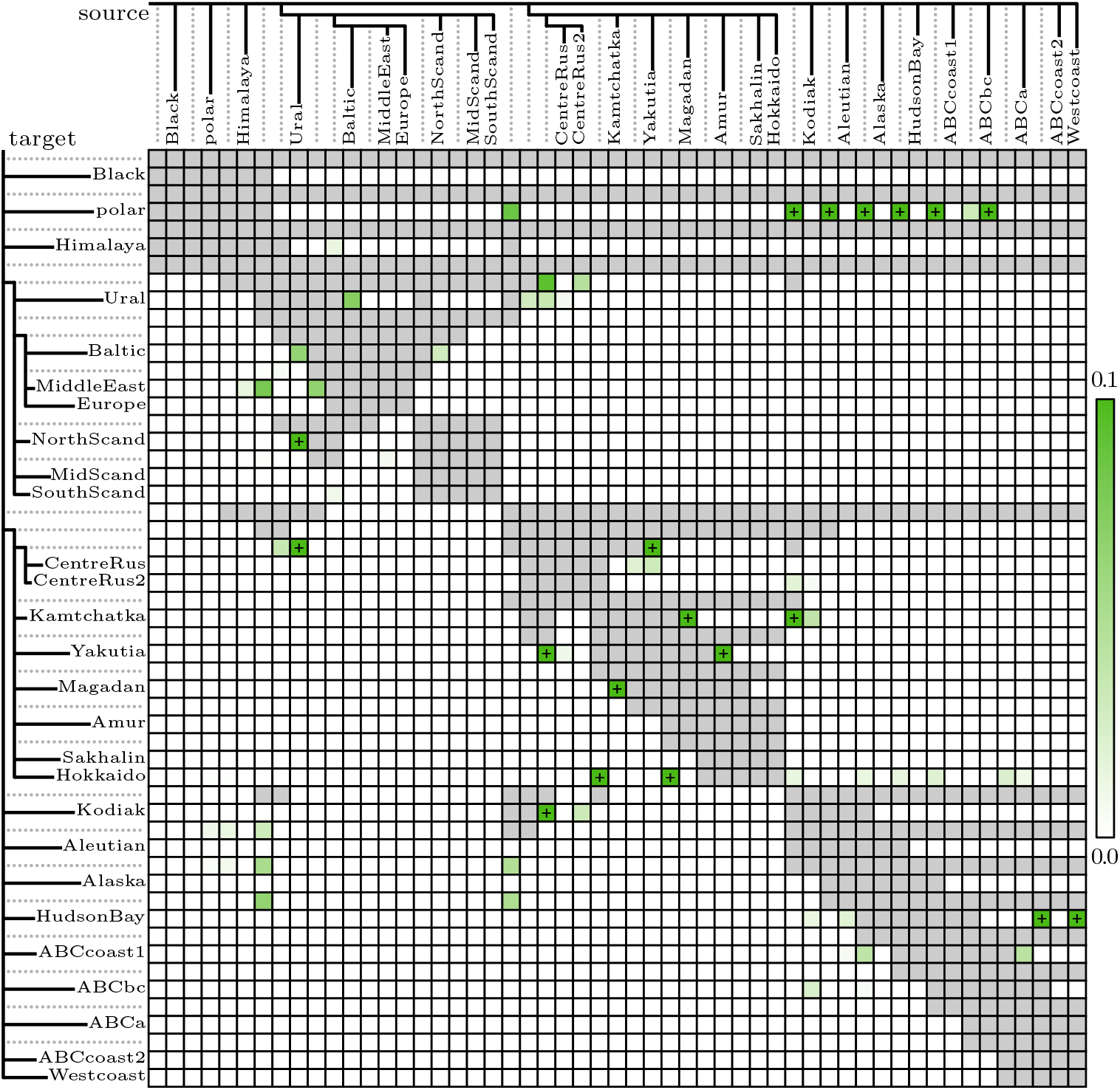
The enhanced branch statistic *f*_*B*_ on 27 bear populations. Grey cell means that the statistic is undefined. The plus symbol means that the value is over 0.1 (the highest value is 0.425). See the main text for discussion.

## Application on brown bear – polar bear gene flow

The brown bears *Ursus arctos* and the polar bears *Ursus maritimus* possess the ability to hybridize and introgress to either parental species by backcrossing. First zoo hybrids were observed in Halle already in 1874 [36] (note that at the time brown bears were not considered a single species). A modern study on accidental zoo hybrids [34] describes both the appearance and the behaviour of the brown bear – polar bear offspring as intermediate between those of the two parental species: light and partially hollow hair, longer neck and tail than what brown bears have, an intermediate amount of plantar fur, hurling toys and using paws to cover their nose while play fighting, resting on the belly with hinds legs stretched back — these are all behaviours not exhibited by brown bears.

In the wild, anecdotal sightings of possible hybrids have been reported earlier, but the first confirmed case was in 2006 [33]. The hybrid was mistaken for a polar bear. The first confirmed case of a second generation hybrid was in 2010 [1]. The introgression was to brown bear — in fact the second generation hybrid was sired by its own maternal grandfather. This was also the case when another four first generation and four second generation hybrids all turned out to be descendants of one polar bear female and two brown bear males [32]. There has not been verified introgression to polar bears in the wild, but nevertheless, the wild hybrids evidently could reach sexual maturity in Arctic conditions, living the polar bear life style. Hybridization in the wild is a rare event; all known instances are from Canadian Northwest Territories where brown bear males occasionally roam far outside their usual range [10]. The consistent pattern where polar bear is the mother and brown bear the father already follows from philopatry of the brown bear females, but it has also been suggested that polar bear males are too timid [7] or too aggressive [37] to mate with brown bears (brown bears and polar bears are known to kill each other [1], but fatal conflict also occurs within species). In any case, the present day regularities regarding either the gene flow direction or the sex bias can only offer limited insight to the dynamics that played out during the deep past when the ranges of the two species fluctuated.

According to maternally inherited mitochondrial DNA polar bears are nested inside the brown bear phylogeny [8, 22]. This was originally not seen as evidence of admixture, because the polar bear was considered a very recent ecological specialization [19] (Kurtén’s oldest sample from Kew Bridge, London, is later suggested to actually be a brown bear [18]). Mitochondrial capture was first proposed in 2011 [11], soon echoed by studies on nuclear genome supporting monophyletic clustering of modern brown bears and cytonuclear discordance [14, 27]. The parsimonious explanation [14, 27] for the discordance is that the original polar bear mitochondrial lineage is lost, and polar bears today (haplogroup 2b) all carry mitochondria they obtained from ABC bears (Admiralty, Baranof and Chichagof islands of Alexander archipelago in Southeast Alaska, where brown bears show the greatest genetic affinity with the polar bears; haplogroup 2a) about 155,000 years ago. Hassanin [17] describes an alternative scenario where the original polar bear mitochondrial line still exists, in which case West European and some Scandinavian bears (haplogroup 1) must also belong to it today if we are to believe that polar bears as a species is at least some half a million years old. According to this scenario, brown bears have captured the polar bear mitochondria at least twice; first about 340,000 years ago in Europe, and again about 155,000 years ago in the ABC islands. The estimates hold up with the most recent data [9, Supplementary figure 3A]. Hassanin points out that temporally the nodes of the mitochondrial phylogenetic tree coincide with glacial stages. In our view, this can just as well be consistent with gene flow from brown bears into polar bears in a scenario where an expanding polar bear population absorbs a brown bear refugium population.

Studies on nuclear genome have not conclusively solved the question of direction. Liu ae al. [23] report that out of the two unidirectional scenarios the IBS tract method [16] favours gene flow from polar bears into brown bears, but that *δ*a*δ*i [13] detects evidence for both directions when bidirectionality is allowed. Cahill et al. [4, 6, 5] argue for gene flow into brown bears alone. Relevant here, the 2015 article [6] is based on an inappropriate use of *f*_4_-ratio statistic where the absence of gene flow into polar bears is deduced by setting both *A* and *B* as polar bears, both *C* and *D* as brown bears, and *E* as the black bear in Figure 2 left panel (see Supplementary figure 20 of [20]). Polar bears form a remarkably homogeneous population, and an ancient gene flow event would affect them all essentially equally — in other words, it’s not reasonable to expect that the gene flow concerns *B* alone, and the *f*_4_-ratio is blind to gene flow into *AB* | *CDE* (right panel). Hence, the role of *A* must be played by the black bears, leaving all *C, D* and *E* to brown bears. This was essentially the insight behind the analysis we reported in Figure 4 panel B of [20], using *f*_2_-statistic fitting and finding evidence for both directions (the sixth leaf in that figure, an ancient polar bear sample, can in this respect be ignored because it just always sticks next to the modern polar bears). The same mistake and the same fix were soon replayed. Wang et al. [38] use the *D*_FOIL_ method [30] on tree *S* = (((1, 2), (3, 4)), 5), where 1 and 2 are brown bears, 3 and 4 are polar bears, and 5 is the black bear. The *D*_FOIL_ method could detect the presence but not the direction of gene flow between American brown bears 1 and all polar bears 34 (the result was however announced unidirectional). Again it was because the homogeneous polar bears form for all practical purposes just one leaf, and again the solution was to introduce a third brown bear leaf. Refining the idea of *D*_FOIL_, in [21] we developed the Δ-statistics that also work on tree *A* = ((((1, 2), 3), 4), 5), and using brown bears as 1, 2 and 3, the polar bear as 4, and the black bear as 5 we could distinguish the two gene flow directions, detecting evidence for both. Other methods have also been applied to the nuclear genome; well fitting admixture graphs agree on brown bear – polar bear gene flow but sometimes favour one direction [9] and sometimes the other [20].

Here, we use the ideas discussed in this note, the enhanced branch statistic *f*_*B*_ and the unrooted *f*_4_-ratio test, to provide evidence for ancient bidirectional gene flow between brown bears and polar bears. We use a recent thinned data set that covers the entire range of brown bears [9], available for download at https://doi.org/10.5061/dryad.qbzkh18n6.

### Using the enhanced branch statistic *f*_*B*_

It’s natural to start with the bird’s-eye view given by the branch statistic. We use the same 27 populations (25 brown bear populations, the black bear and the polar bear) as de Jong et al. [9], organized into the same backbone tree as was used in the *TreeMix* analysis on Figure 4 panel g. As reverse gene flow causes false positive signals when *n* = 1, and we seek to illuminate the brown bear – polar bear gene flow, we this time required *n* ≥ 2. The results are depicted in Figure 6, and the corresponding analysis using Malinsky’s *f*_*b*_ can be found in Supplementary figure 3.

The most striking result is the signal for gene flow from brown bears into polar bears. Because the event *s* → *t* will make *f*_*B*_ implicate all gene flow events *s*_1_ → *t* where *s*_1_ is an ancestor of *s* (see discussion on Figure 4 lower right panel), the parsimonious explanation is gene flow from ABCbc into polar. The way *f*_*B*_ is defined means the signal is persistent with respect to the choice of 𝒟 and ℰ, and according to Figure 2 right panel, the only gene flow event that alone could make the *f*_4_-ratio positive independently of the choice of *D* and *E* is *C* → *B*. The consistency with 𝒟 also rules out some more complex scenarios, such as the combination of reverse gene flow *B* → *C* and subsequent *C* → *D*.

The opposite direction is only supported by weak signals, most notably of gene flow from polar into the ancestors of all American brown bears except Kodiak (*f*_*B*_ = 0.00892). Because of the dense sampling among brown bears, the branch statistics *f*_*b*_ and *f*_*B*_ are not ideal for assessing the gene flow form polar bears into brown bears. When for example 𝒞 is polar and ℬ is ABCbc, the statistic *f*_*B*_ attempts to isolate only the gene flow that is private to ABCbc. As 𝒜 is now fixed to {ABCa, ABCcoast2, Westcoast}, American brown bears admixing with both polar bears and each other makes the *f*_4_-ratio erratic, attaining even predominantly negative values when ℰ is Kodiak (note that we also detect gene flow within brown bears from Kodiak into ABCbc, and that according to Figure 2 right panel the event *E* → *B* has a negative effect on the *f*_4_-ratio) or the clade of Far Eastern brown bears (Supplementary table 1). Consequently *f*_*B*_ = 0. For the general question of whether American brown bears have polar bear ancestry, it is better to reduce the resolution by removing or pooling some of the American samples and thereby pushing the set 𝒜 further back. Already removing ABCcoast1, ABCa, ABCcoast2 and Westcoast makes the *f*_4_-ratio stable, indicating gene flow into the ABCbc lineage after splitting from the HudsonBay lineage (Supplementary table 2).

The various gene flow events detected within the brown bear populations suggest that a tree is a poor model of the brown bear diversity. Roughly, Ural and the central Russian samples connect Eurasia through paths outside the tree, and Kodiak acts as a bridge between Eurasia and America. This is in line with de Jong et al. [9] presenting several conflicting perspectives to autosomal brown bear structure: the backbone used by *TreeMix* (Figure 4 panel g) and by us, the *bioNJ dendrogram* (Figure 1 panel f, Figure 2 panel a), the *Neighbor-Net algorithm network* (Figure 4 panel d), the *ASTRAL supertree* (Figure 5 panel b), and the *neighbour joining tree* (Figure 5 panel d). According to most models, the European cluster is basal compared to Far Eastern and American clusters, but even this is contested by the *ASTRAL supertree*.

### Estimating the admixture proportions

To estimate gene flow proportions, we will use *A* = ((((1, 2), 3), 4), 5), where 5 is Black, 4 is polar and 1–3 are brown bear populations. Including one Eurasian sample as 3 and two American samples as 1 and 2 minimizes the bias from within brown bear gene flow; this way we sidestep the basal ambiguity between the European, Far Eastern and American clusters. Fixing 2 as ABCbc, Figure 7 visualizes the unidirectional *f*_4_-ratio estimates for all choices of 1 and 3.

**Figure 7.**
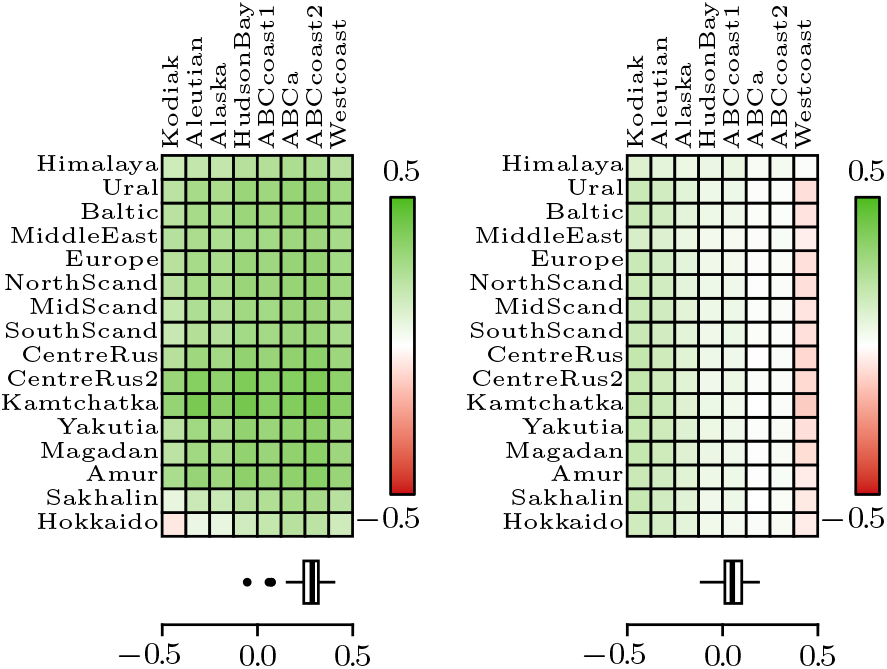
Left: unidirectional *f*_4_-ratio estimates for proportion of gene flow form brown bears into polar bears. Here *A* = Black, *B* = polar, *C* = ABCbc, *D* is an American brown bear (columns), and *E* is an Eurasian brown bear (rows). Otherwise consistent effect is less pronounced in the island populations of Sakhalin and Hokkaido, in particular if America is represented by the Kodiak island (−0.054, SE = 0.079). Right: unidirectional *f*_4_-ratio estimates for proportion of gene flow form polar bears into brown bears. Here *A* is an American brown bear (columns), *B* = ABCbc, *C* = polar, *D* = Black, and *E* is an Eurasian brown bear (rows). The gradient with respect to the American representative reflects distance between *A* and *B*, as the *f*_4_-ratio only detects gene flow that is private to *B*, not affecting *A*. With Westcoast the estimates even become negative (−0.119, SE = 0.014 when *E* is Kamtchatka).

Based on Figure 7 left panel, the gene flow from brown bears into polar bears is well supported. The estimated proportion is consistent and largely independent of the choices of *D* and *E*, with the exception that using the island samples from Sakhalin and Hokkaido appears to dilute the signal (the negative value is not statistically significant). This could be due to island populations partly representing lost population structure [9], although we don’t understand the exact mechanism. Independence of the choice of the Eurasian sample *E* gives confidence that gene flow from extinct cave bears *Ursus spelaeus* [2] is not confounding the results.

The right panel suggests gene flow from polar bears into brown bears. The estimates are virtually independent of the choice of the Eurasian sample *E*, which again rules out the influence of cave bear admixture. This time *D* is held constant at Black, which raises the possibility of a *D*-specific bias. These could include *D* → *B* or *D* → *A*, which are contradicted by ABBA-BABA-statistics showing that American brown bears have more affinity to polar bears than Eurasian brown bears do [4], or *B* → *C* and subsequent *C* → *D*, which we dismiss out of hand because it relies of gene flow from polar bears into black bears. We deduce *C* → *B*, which is reflected in the gradient with respect to the choice of the American sample *A*. The simplest explanation for the significantly negative values when *A* = Westcoast is ABCbc not actually being the brown bear population with the most polar bear ancestry; we will return to this point momentarily.

For bidirectional estimates we choose representatives 1 = Aleutian because it’s distant from ABCbc and because Kodiak is heavily involved in within brown bear gene flow (see Figure 6), and 3 = Europe because it has the most individuals among the Eurasian samples that are not the source or the target of within brown bear gene flow (Europe, MidScand, Sakhalin, see Figure 6). The second row in Table 1 has the appropriate formulae, significance was assessed using the block jackknife. For the proportion of gene flow from brown bears into polar bears we get 0.243, SE = 0.035 (unidirectional estimate 0.271, SE = 0.037). For the proportion of gene flow from polar bears into brown bears we get 0.105, SE = 0.009 (unidirectional estimate 0.138, SE = 0.011). Interestingly, replacing ABCbc with Westcoast gives overall higher estimates: from brown bears into polar bears 0.260, SE = 0.037 (unidirectional estimate 0.304, SE = 0.041) and from polar bears into brown bears 0.144, SE = 0.011 (unidirectional estimate 0.195, SE = 0.012). Here it looks like the ABC bears might not be the population most admixed with polar bears. On the other hand, *f*_4_(ABCbc, Europe; polar, Black) = 0.00672 (SE = 0.0003) and *f*_4_(Westcoast, Europe; polar, Black) = 0.00346 (SE = 0.0003), but we should keep in mind that a simple *f*_4_-statistic (like the ABBA-BABA-statistic) is not directly targeting the gene flow proportions but depends on timing and branch lengths as well.

## Acknowledgements

This work was supported by Academy of Finland Center of Excellence in Tree Biology (TreeBio AoF CoE 346139). We thank Benjamin C Haller for teaching at a *SLiM* workshop in summer 2025 in Helsinki, as well as the local organizers.

## Supplementary material

**Supplementary figure 1.**
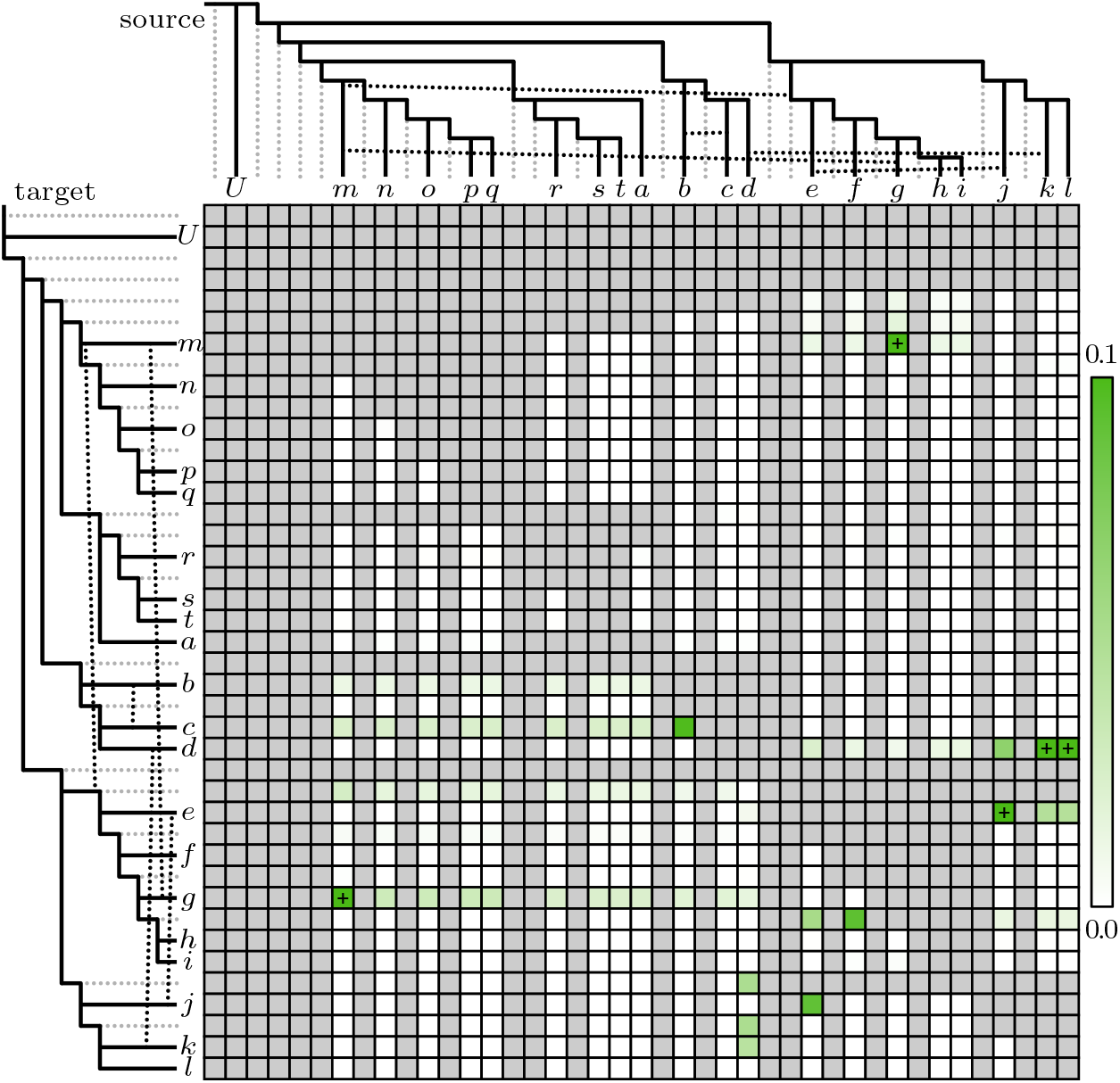
The left panel of Figure 5 embedded on a table where the source of gene flow is allowed to be an inner branch, which Malinsky’s branch statistic *f*_*b*_ doesn’t permit. Grey cell means that the statistic is undefined. The plus symbol means that the value is over 0.1 (the highest value is 0.149). The purpose of this figure is only to contrast it to Supplementary figure 2 of the more liberal enhanced branch statistic *f*_*B*_.

**Supplementary figure 2.**
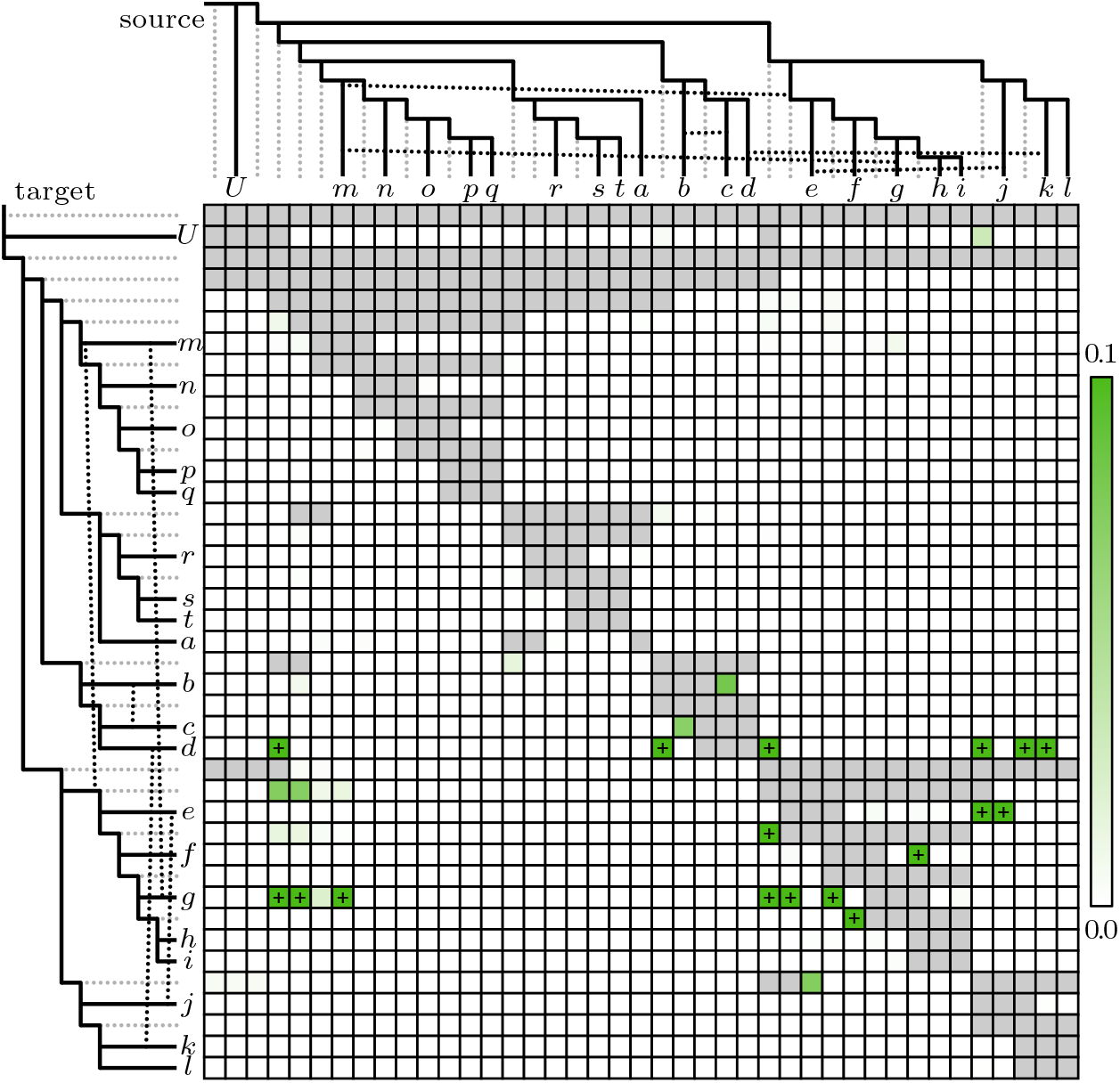
The right panel of Figure 5 expanded on a table where the source of gene flow is allowed to be an inner branch. Grey cell means that the statistic is undefined. The plus symbol means that the value is over 0.1 (the highest value is 0.192). Compared to Supplementary figure 1 we can appreciate the broader definition of *f*_*B*_. As discussed in relation to Figure 4 lower right panel, pinpointing the source of gene flow is hindered by the fact that *f*_*B*_ tends to implicate the parental (but not descendant or sister) branches of the true source of gene flow as a source as well. When several sources are indicated, it’s therefore advisable to trust the signal with largest *n*. For example, there results suggest *k* → *d* (*n* = 6), *kl* → *d* (*n* = 5), *jkl* → *d* (*n* = 4), *efghijkl* → *d* (*n* = 3), *mnopqrstabcd* → *d* (*n* = 2) and *bcd* → *d* (*n* = 1), out of which only the first one is reality. Note that the latter three false source branches were only “parental” to *k* in an unrooted sense. As in Figure 5, when *n* = 1 the false positives tend to occur more. Apart from these two easily understood sources of wrong signals, we only observe faint *g* → *m, mnopqrsta* → *fghi* and *mnopqrstabcd* → *fghi* that are not true.

**Supplementary figure 3.**
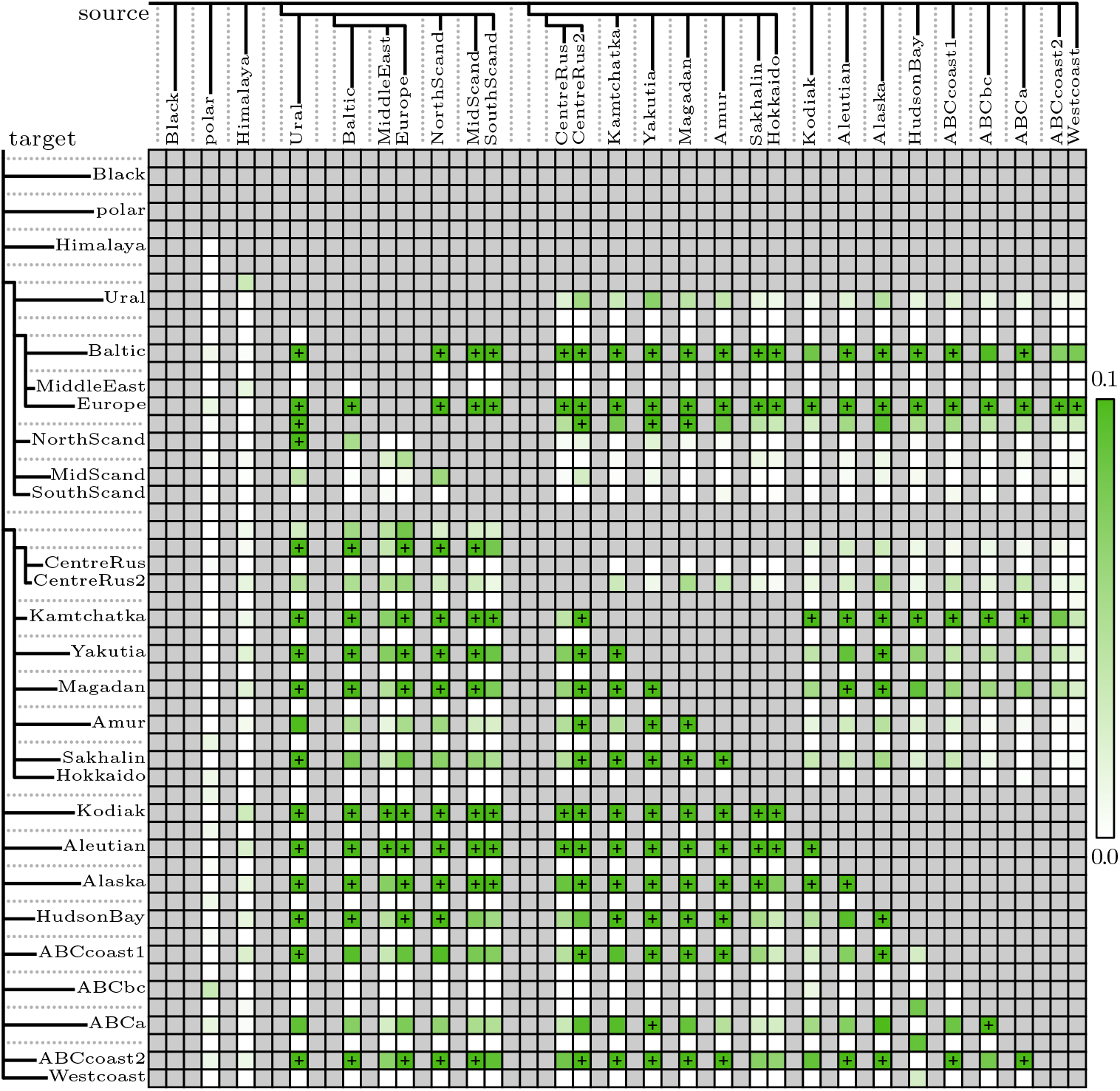
Using Malinsky’s *f*_*b*_ instead of the enchanced *f*_*B*_ in Figure 6. Gene flow from brown bears into polar bears cannot be assessed because of the built-in restriction of requiring there exists an outgroup and that the outgroup is not in 𝒜. Signal of gene flow from polar bears into brown bears is more pronounced than in Figure 6, but it could just as well be due to reverse gene flow, see Figure 4 top right panel. A lot of the within brown bear gene flow signal is likely false positives due to difficulties in detecting the actual source of gene flow, as was illustrated in Figure 4 bottom right panel and its explanation.

**Supplementary table 1.** An enhanced statistic branch *f*_*B*_ dissected. Values of 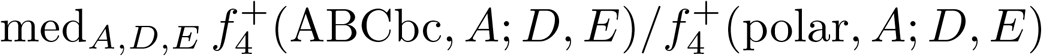 for different sets 𝒟, *ε*. Here “European” consists of populations Ural, Baltic, MiddleEast, Europe, NorthScand, MidScand and SouthScand, and “Far Eastern” consists of populations CentreRus, CentreRus2, Kamtchatka, Yakutia, Magadan, Amur, Sakhalin and Hokkaido. The labels are not faithful to geography, but follow the clades in the backbone phylogeny. In this resolution the statistic is unstable as the set 𝒜 = {ABCa, ABCcoast2, Westcoast} is close and mixing with ABCbc, as well as carrying its own polar bear ancestry. Gene flow from Kodiak into ABCbc also has an effect. Some of the lower medians become zero, and so does the branch statistic *f*_*B*_.

| | | $\mathcal{E}$ | | | | | | | | |
| --- | --- | --- | --- | --- | --- | --- | --- | --- | --- | --- |
|  |  | Black | Himalaya | European | Far Eastern | Kodiak | Aleutian | Alaska | HudsonBay | ABCcoast1 |
| $\mathcal{D}$ | polar | 0.034 | 0.034 | 0.021 | 0.020 | 0.017 | 0.019 | 0.030 | 0.043 | 0.029 |
|  | Black |  | 0.025 | 0.007 | 0.005 | 0.000 | 0.007 | 0.022 | 0.063 | 0.031 |
|  | Himalaya |  |  | 0.000 | 0.000 | 0.000 | 0.000 | 0.018 | 0.108 | 0.027 |
|  | European |  |  |  | 0.000 | 0.000 | 0.006 | 0.101 | 0.211 | 0.107 |
|  | Far Eastern |  |  |  |  | 0.000 | 0.000 | 0.119 | 0.238 | 0.105 |
|  | Kodiak |  |  |  |  |  | 0.609 | 0.537 | 0.593 | 0.259 |
|  | Aleutian |  |  |  |  |  |  | 0.455 | 0.585 | 0.232 |
|  | Alaska |  |  |  |  |  |  |  | 0.704 | 0.007 |
|  | HudsonBay |  |  |  |  |  |  |  |  | 0.000 |
| med |  | 0.034 | 0.025 | 0.007 | 0.000 | 0.000 | 0.006 | 0.101 | 0.211 | 0.031 |

**Supplementary table 2.** An enhanced statistic branch *f*_*B*_ dissected. Values of 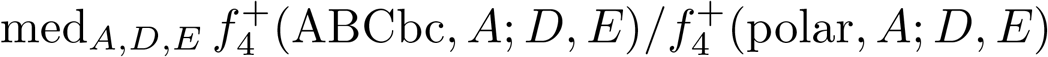 for different sets 𝒟, ℰ and ABCcoast1, ABCa, ABCcoas2 and Westcoast removed. Here “European” consists of populations Ural, Baltic, MiddleEast, Europe, NorthScand, MidScand and SouthScand, and “Far Eastern” consists of populations CentreRus, CentreRus2, Kamtchatka, Yakutia, Magadan, Amur, Sakhalin and Hokkaido. The labels are not faithful to geography, but follow the clades in the backbone phylogeny. In this resolution the statistic exhibits signs of gene flow from polar into ABCbc, obtaining a positive value of 0.048. With Kodiak as 𝒟 and Aleutian as ℰ the ratio is not defined as the denominator is never positive.

| | | $\mathcal{E}$ | | | | | | |
| --- | --- | --- | --- | --- | --- | --- | --- | --- |
| $\mathcal{D}$ | | Black | Himalaya | European | Far Eastern | Kodiak | Aleutian | Alaska |
|  | polar | 0.048 | 0.050 | 0.049 | 0.049 | 0.050 | 0.050 | 0.051 |
|  | Black |  | 0.059 | 0.053 | 0.052 | 0.055 | 0.054 | 0.060 |
|  | Himalaya |  |  | 0.027 | 0.026 | 0.046 | 0.043 | 0.061 |
|  | European |  |  |  | 0.093 | 0.066 | 0.059 | 0.088 |
|  | Far Eastern |  |  |  |  | 0.056 | 0.049 | 0.086 |
|  | Kodiak |  |  |  |  |  |  | 0.159 |
|  | Aleutian |  |  |  |  |  |  | 0.254 |
|  | med | 0.048 | 0.050 | 0.049 | 0.049 | 0.055 | 0.050 | 0.086 |

## Notes

https://github.com/KalleLeppala/rootless

### Competing Interest Statement

The authors have declared no competing interest.

